# Improving the specificity of adenine base editor using high-fidelity Cas9

**DOI:** 10.1101/712109

**Authors:** Ruisha Hong, Sidi Ma, Feng Wang

**Affiliations:** State Key Laboratory of Biocontrol, School of Life Sciences, Sun Yat-sen University, Guangzhou 510275, China

**Keywords:** CRISPR/Cas9, high-fidelity ABE, off-target activity, pathogenic SNPs

## Abstract

Adenine base editor (ABE) mediates the conversion of A to G in genomic DNA. In human, approximately 47.8% of known pathogenic SNPs can be corrected by A to G conversion, indicating that ABE have tremendous potential in gene therapy. However, the off-target activity of ABE limits its clinical application. ABE off-target activity in DNA is depended on the bonding of *Streptococcus pyogenes* Cas9 (SpCas9) on off-target sites [1, 2]. Therefore, using high-fidelity Cas9 should be able to improve the specificity of ABE. Based on this, we replaced the wild-type SpCas9 in ABE7.10 with four high-fidelity Cas9s to improve its specificity. The analysis of target deep sequencing data demonstrate that the specificity of e-ABE is substantially improved compared to conventional ABE7.10 in four test sites. But the broad editing window of ABE hampers its application, ABE needs to be optimized to get variants with narrow editing window.

## 1. Introduction

The CRISPR (clustered regularly interspaced short palindromic repeats)-Cas (CRISPR-associated proteins) is an adaptive immune system widely existing in bacteria and archaea, which can cleavage invading foreign nucleic acids directed by short guide RNAs [3, 4]. The system has been engineered as gene editing tools, including a single guide RNA (sgRNA) and Cas9 nucleases [5, 6]. The sgRNA-Cas9 complex cleavages target DNA sequence based on two conditions: (1) compatibility of the protospacer adjacent motif (PAM) with the PAM-interacting domain of Cas9, and (2) complementary of the sgRNA sequence with the target site. However, the nuclease activity of Cas9 may also be triggered when there are mismatches between sgRNA and off-target site [7, 8]. These off-target effects limit its application in genome-editing, especially in clinical.

Several strategies have be used to reduce the off-target activity of Cas9, including reducing the amount of active Cas9 [9], using paired nickases Cas9 (nCas9) [10, 11], fusing Cas9 with specific DNA-binding proteins [12, 13], engineering of the sgRNAs (truncated sgRNAs [14] and hairpin sgRNAs [15]) and so on. In addition, several high-fedility Cas9 variants have been developed, such as enhanced specificity Cas9 (eSpCas9) [16], Cas9-high fidelity (Cas9-HF) [17], evoCas9 [18], HypaCas9 [19] and sniper-Cas9 [20].

Based on CRISPR/Cas9, single base editing system has been developed, including cytosine base editors (CBEs) [21] and adenosine base editors (ABEs) [2]. CBEs enable C•G to T•A base pair conversion, while ABEs enable A•T to G•C base pair conversion, at a target genomic locus without inducing double strand breaks. ABEs consist of two parts, sgRNA and fusion protein of adenosine deaminase TadA and nCas9. Locating in a target site guided by sgRNA, TadA catalyzes base A deamination to I (inosine) on the non-complementary strand, meanwhile nCas9 nick the complementary strand. Because base I can pair with C, A•T convert to G•C after replication.

In human, about 47.8% of known pathogenic single nucleotide polymorphisms (SNPs) can be corrected by conversion of A•T to G•C (Fig. 3A), indicating ABEs have tremendous potential in gene therapy [2]. However, ABEs can tolerate mismatch between sgRNA and target sequence, inducing off-target effects [22], which limit its application especially in clinical. The using of high-fidelity Cas9 variants in ABE have not been reported, except sniper-Cas9. But sniper-ABE also have off-target activity on some sites [23]. Because the high-fidelity Cas9 variants not always have same off-target sites [18], we need various high-fidelity ABEs. Here, we replaced the wild type SpCas9 in ABE7.10 with four high-fidelity Cas9s (eSpCas9, Cas9-HF, evoCas9 and hypaCas9), named e-ABE7.10, HF-ABE7.10, evo-ABE7.10 and Hypa-ABE7.10, in order to enhance the specificity of ABE.

## 2. Materials and methods

### 2.1. Plasmid construction

The conventional ABE7.10 plasmid (pCMV-ABE7.10; Addgene, #102919) was purchased from Addgene, besides we optimized its codon according previous report [24].The sgRNA plasmid pUC57-Cas9-gRNA was synthesized by GENEWIZ. The pCMV-eABE7.10, pCMV-HypaABE7.10 and pCMV-evoABE7.10 plasmids were constructed through site directed mutagenesis by PCR on pCMV-ABE7.10 plasmid backbone, and mutated sites are shown in Fig. 1A. The pCMV-HFABE7.10 plasmid was constructed through Gibson assembly of necessary sequence (synthesized by GENEWIZ) into the pCMV-ABE7.10. Primers used for gRNA inserting into the pUC57-Cas9-gRNA plasmid are shown in Supplementary Table 3. All plasmids were confirmed by Sanger sequencing.

**Fig. 1.**
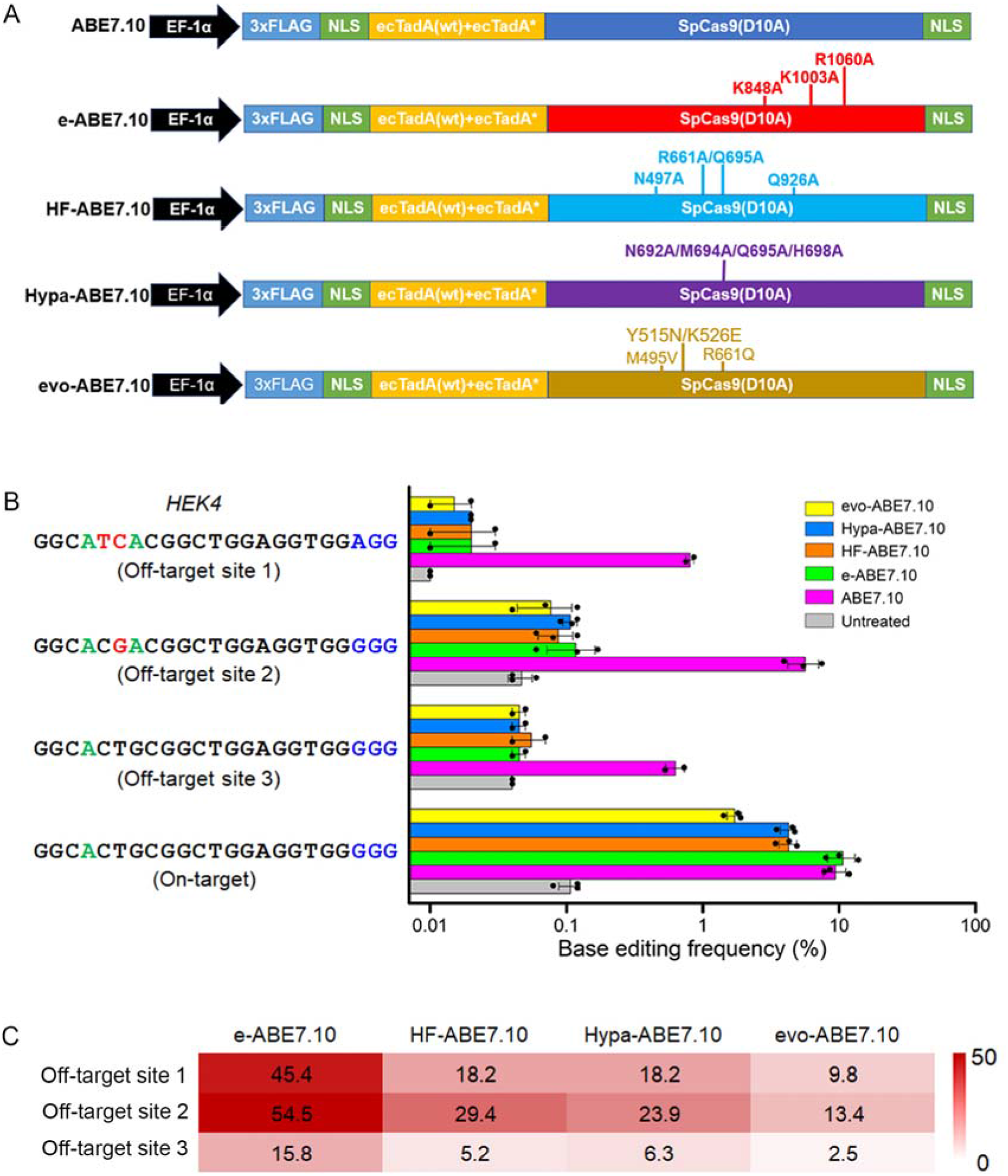
Construction of the high-fidelity ABEs and its base editing analysis. (A) Schematic representation of ABE.10, e-ABE7.10, HF-ABE7.10, evo-ABE7.10 and Hypa-ABE7.10. (B) Base editing efficiencies of ABEs measured by targeted deep sequencing at HEK4 on- and off-target sites in HEK293T cell. Mismatched bases, edited bases and PAM sequences are shown in red, green and blue, respectively. (C) Specificity ratios were shown by the heatmap, calculated by the formula: (high-fidelity ABE7.10 on-target frequency/off-target frequency)/(ABE7.10 on-target frequency/off-target frequency). Means ± SD were from two or three independent experiments.

### 2.2. Human cell culture and transfections

HEK293T cells were cultured and passaged in Dulbecco’s modified Eagle’s medium (DMEM, Gibco) supplemented with 10% (v/v) fetal bovine serum (FBS, Gibco) and 1% (v/v) penicillin-streptomycin (Gibco). Cell lines were maintained at 37 °C with 5% CO_2_. Plasmids for mammalian cell transfections were prepared using an Endo-free Plasmid Mini Kit II (OMEGA). HEK293T (1.5× 10^5^ cells) were seeded into 24-well Poly-d-Lysine-coated plates (Corning) in the absence of antibiotic. After12-15 hours plating, 750 ng ABE plasmid and 250 ng gRNA plasmid were co-transfected into the cells with Lipofectamine 3000 according to the manufacturer’s protocol. Genomic DNA was extracted 5 days after transfection by Tissue DNA Kit (OMEGA).

### 2.3. DNA amplification and deep sequencing

Primers used for on- and off-target sites PCR amplification were listed in Supplementary Table 3. PCR was performed using Q5 Host Start High- Fidelity 2× Master Mix (NEB), and 150 ng Genomic DNA was chosen as the template. The program of PCR: 98°C for 30 s, then 35 cycles of (98°C for 10 s, 67°C for 10 s, and 72°C for 10 s), followed by a final extension of 2 min at 72°C. PCR products were purified by electrophoresis and deep sequenced using the HiSeq 2×150bp sequencing system (Illumina) as paired-end 150 reads at GENEWIZ.

### 2.4. Data analysis

Because the length of reads didn’t cover the whole fragments sequenced. For data sequenced by NGS platforms, we used FLASH [25] to merge the pair-ends sequence reads to do the following process. The nucleotide constitution analysis were performed with python referencing the python script in a previous paper [2]. Some adjustments were done to fit our experiment requirements.

For editing site analysis, we used the ClinVar data with “fileData = 2019-06-18”. We chose data in VCF format with the INFO of “CLNSIG=Pathogenic”, i.e. clinical significance for this single variant was pathogenic. Using these data could assess candidates for base editing therapy. The potential window size varied between 1 and 8, and latent editing windows were located in fragment between third base and tenth base. The human genome analyzed is Genome Reference Consortium Human Build 38 (GRCh38). The data and scripts are available from the authors upon reasonable request.

## 3. Results

We selected the well validated site *HEK4* [2] to test the specificity of the four ABEs. On the three known off-target sites, the specificity of four ABEs increased to 2.5 - 54.5 times, compared with ABE7.10 (Fig. 1 B, C). The on-target editing activity of e-ABE7.10 is comparable to ABE7.10, but three others have varying degrees of reduction compared to ABE7.10. Therefore, we chose e-ABE7.10 for further testing on the other known sites (*HBG2*, *HPRT* and *VEGFA3* [22, 23]). As shown in Fig. 2, the specificity ratios are increased to 1.1 - 44.2 times, except *HPRT* off-target site 2 and 3. This is due to that both e-ABE7.10 and ABE7.10 have no observable base editing at the *HPRT* off-target site2 and 3, but ABE7.10 have a higher base editing activity at *HPRT* on-target than e-ABE7.10. Compared with ABE7.10, the on-target base editing activity of e-ABE7.10 maintained the same level at site *VEGFA3*, but it have some decrease at sites *HBG2* and *HPRT*. Collectively, e-ABE7.10 has a high specificity in a site dependent method.

**Fig. 2.**
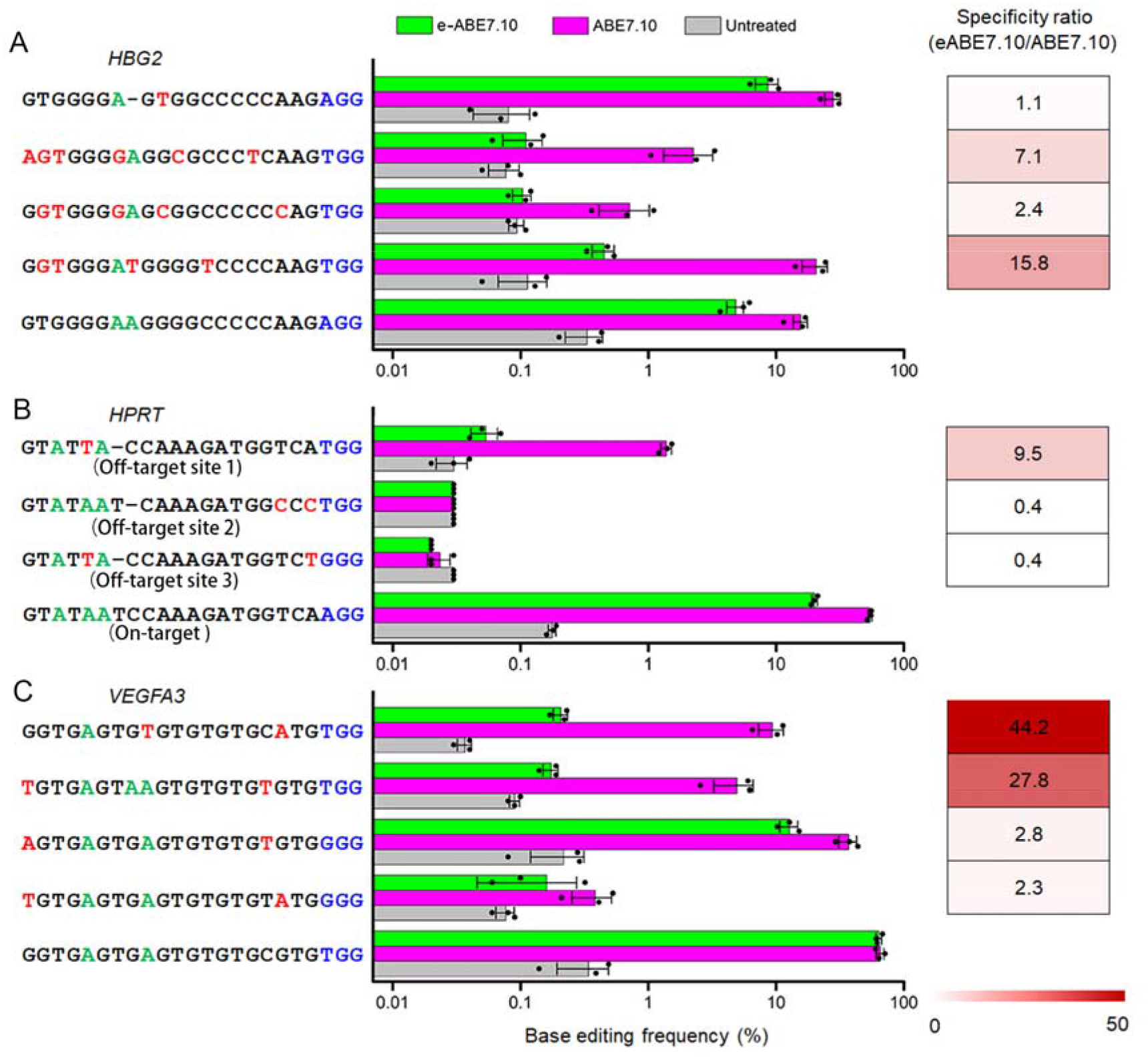
Base editing analysis of eABE7.10. Base editing efficiencies of eABE7.10 and ABE7.10 measured by targeted deep sequencing at *HBG2* (A), *HPRT* (B) and *VEGFA3* (C) on- and off-target sites in HEK293T cells. Mismatched bases, edited bases and PAM sequences are shown in red, green and blue, respectively. Dashes indicate presumed RNA bulges. Specificity ratios were shown by the heatmap, calculated by the formula in Fig. 1. Means ± SD were from three independent experiments.

**Fig. 3.**
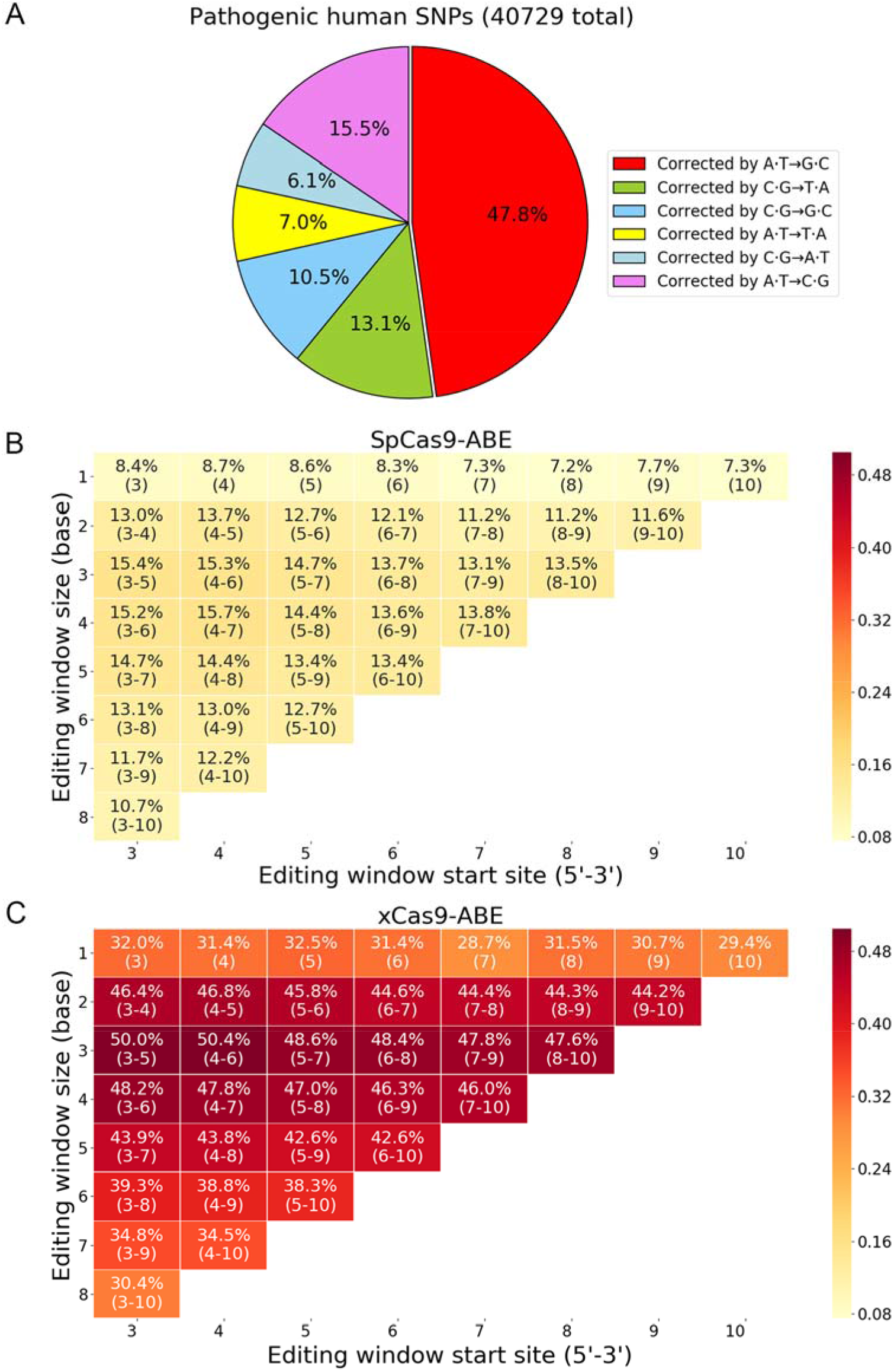
Pathogenic SNPs can be corrected using ABEs. (A) Base pair changes need to correct pathogenic SNPs in the ClinVar Datebase[26]. The percent of pathogenic SNPs (G to A) can be corrected by SpCas9-ABE (B) and xCas9-ABE (C). The assumed ABEs have different editing window.

We find that both e-ABE7.10 and ABE7.10 also have base editing activity at the position 3 and 9 of protospacer, consistent with previous reports [2]. The broad editing windows hamper its application in clinical due to it may contain more than one A in the window. The canonical ABE7.10 original editing window is between third base and tenth base from 5’ end to 3’end [2]. Many adenines in this fragment are edited by ABE leading to wrong editing. One solution to this defect is shrinking the editing windows from 8 bases fragments to smaller fragments. But we have no idea where the new editing window will arise in the editing window, every position in the editing window is possible. As the windows length and location relative to 5’ end changes, SNPs’ editable situations also alter much.

There are two factors to determine whether one SNP can be corrected using ABE without wrong editing in editing window. Firstly, the longer new editing windows are, the more probably additional non-target adenines will arise in editing windows which is not acceptable for clinical use. Secondly, the longer editing windows are, the more likely the PAMs will find, because target adenine can change its location in editing windows allowing find PAM in other places. For example, for window size of 8, a pathogenic SNP can choose 8 possible distance between target site and PAM sequences. But for window size of 1, the position of target nucleotide is deterministic and the PAM position is exclusive. These two factors are antagonist, so we analysis all the potential target inducing human disease stored in ClinVar datebase [26]. When the editing window is in positions 3-9, 11.7% and 34.8% of pathogenic G to A SNPs can be corrected by spCas9-ABE and xCas9-ABE, respectively (Fig. 3). The xCas9 has a broad PAM range (5′-NG, 5′-GAA, 5′-GAT, and 5′-CAA) [27]. If the editing window is in positions 4-8, the percent will increase to 14.4% and 43.8%. We analyzed the possible ABEs with different editing window 3-10, indicating that the ABE with editing window 4-7 have largest target number in SpCas9-ABE but the window is 4-6 in xCas9-ABE. To summarize, we should engineer ABE variants with different editing window, especially with window 4-6 in xCas9-ABE.

## 4. Discussion

To reduce or avoid ABE off-target activity, we replaced ABE’s original Cas9 with four high-fidelity Cas9. At different loci, e-ABE showed 1.1 to 54.5 times improvement in specificity ratios, compared with ABE7.10. eSpCas9 consists three neutral alanine substitution at positively charged residues within the nontarget strand groove, requiring more stringent Watson-Crick base pairing between sgRNA and the target DNA strand for the single strand formation of nontarget DNA strand [16]. The single DNA strand is needed for base A deamination by ABE [2]. It should also be able to improve the specificity of CBE. Extended sgRNA can reduce ABE off-target activity[23], it may have the same effect in e-ABE7.10, needing further test.

According to recent reports, the off-target sites of ABE is fewer in vitro analysis by whole-genome sequencing [22], and in vivo, the same results were observed in rice [1] and mouse embryos [28]. So we only chose the top three or four off-target sites with relatively strong off-target activity to test the specificity of ABEs. But we should perform more comprehensive test, in order to get more convincing results. At the same time, we realized that the ABE with broad editing window will hamper its application in clinical (Fig. 3), but we do not have an ABE with narrow editing window by now. Engineering ABE variants with narrow window is a more important task. There is still a long way to go and more manpower and material resources should be invested in the field.

## Conflicts of interest

The authors declare no conflict of interest.

## Acknowledgements

We thank Dr. Xionglei He for his advices on experiments.

