## Supplemental Table 1-3 for "Improving the specificity of adenine base editor using high-fidelity Cas9"

**Supplementary materials**

**Supplementary Table 1.** Target deep sequencing results at the *HEK4* on target in HEK293T cells treated with ABE7.10, e-ABE7.10, HF-ABE7.10, Hypa-ABE7.10 and evo-ABE7.10. One arbitrary replicate is shown.

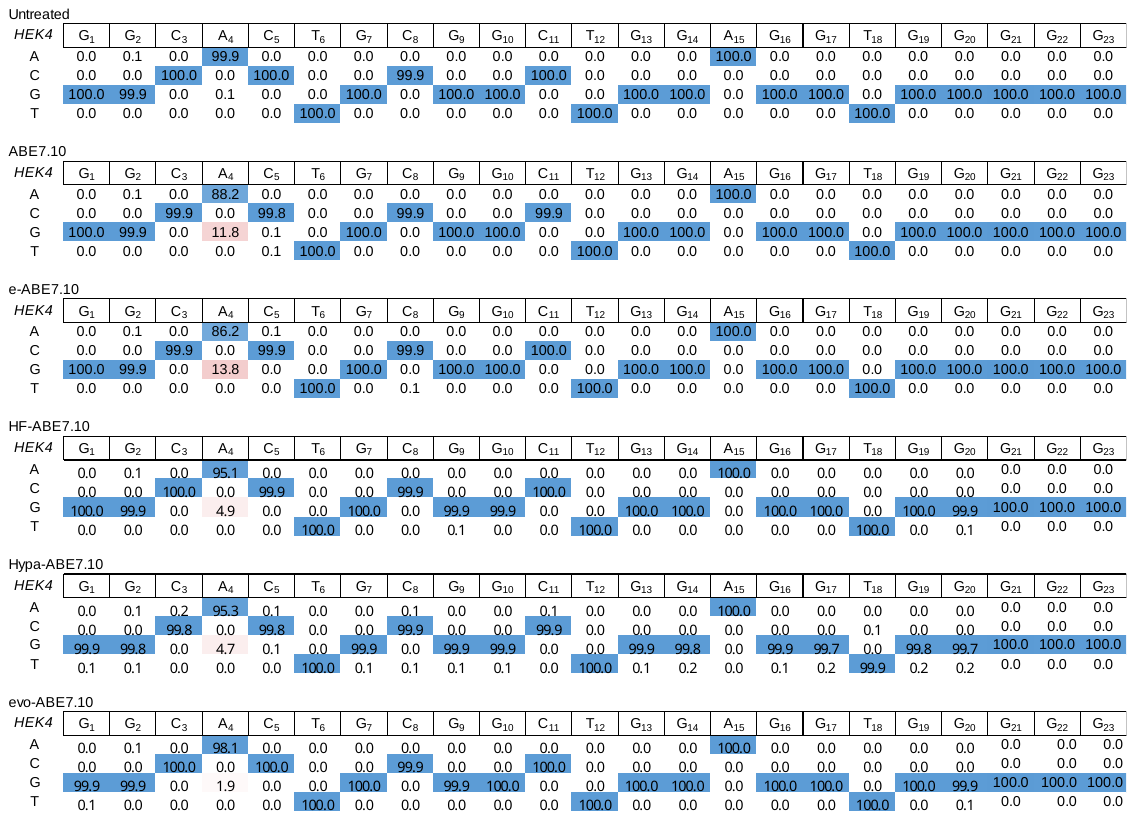

**Supplementary Table 2.** Target deep sequencing results at the sites *HBG2*, *HPRT* and *VEGFA3* on target in HEK293T cells treated with ABE7.10 and e-ABE7.10. One arbitrary replicate is shown.

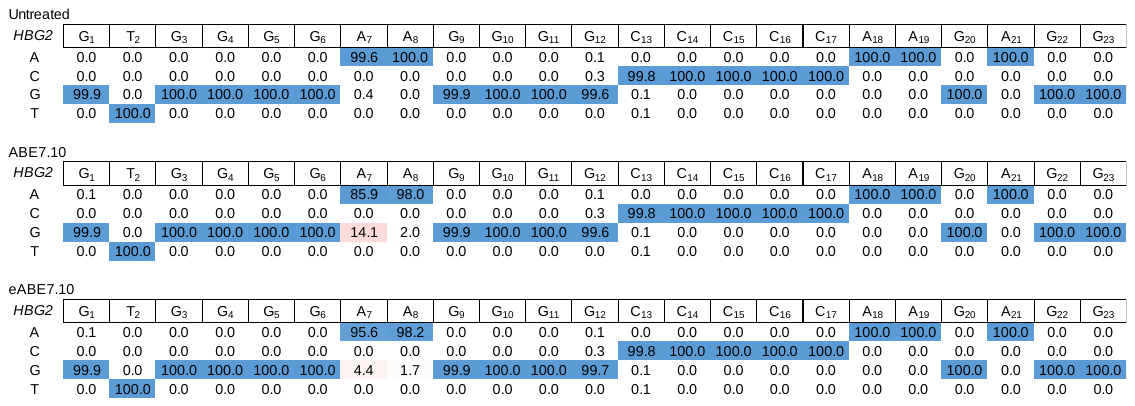

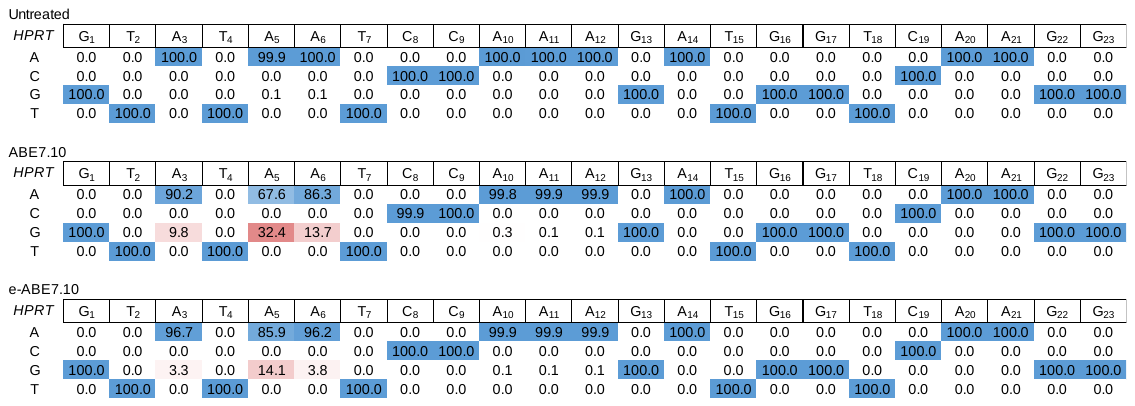

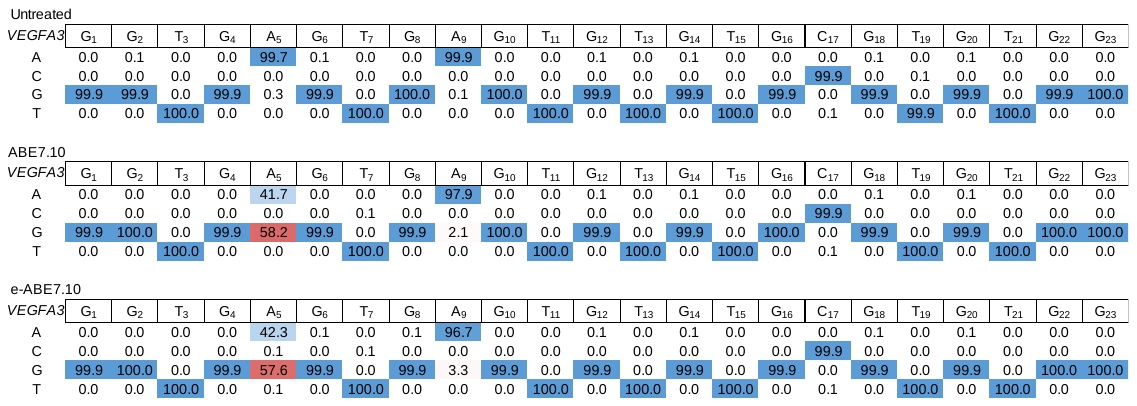

**Supplementary table 3.** Primers used for generating sgRNA plasmids and genomic DNA amplification. Barcode sequences are shown in red.

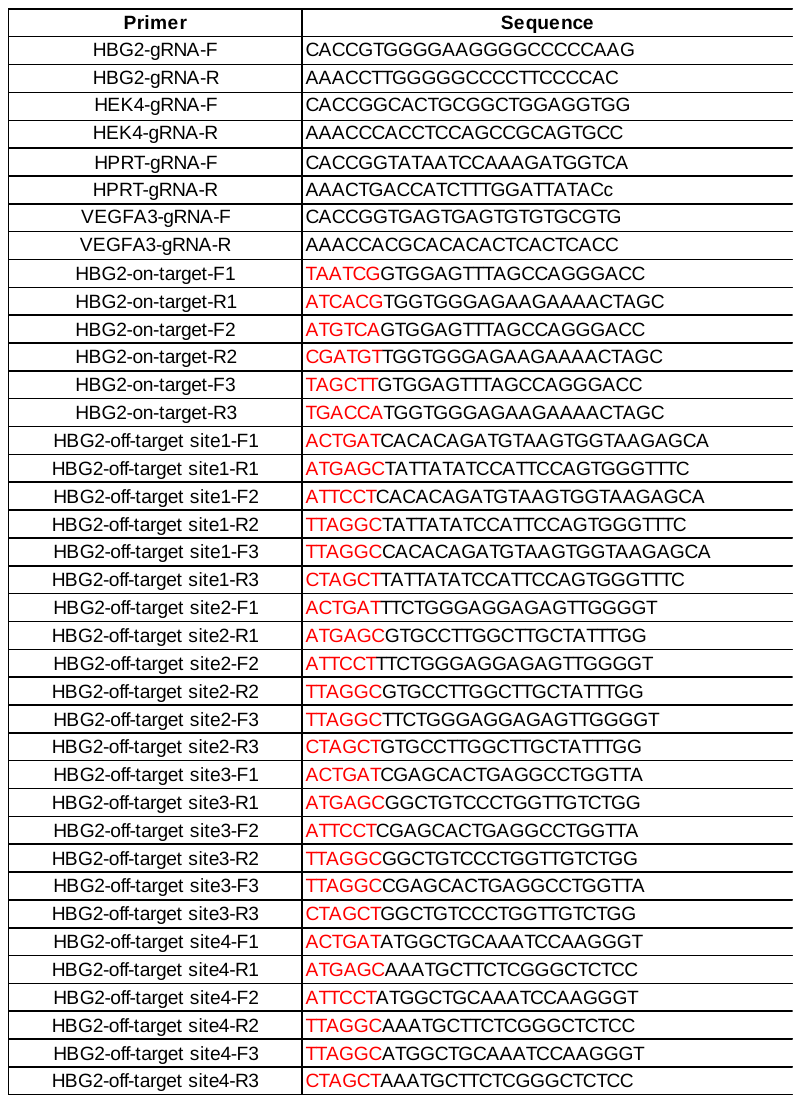

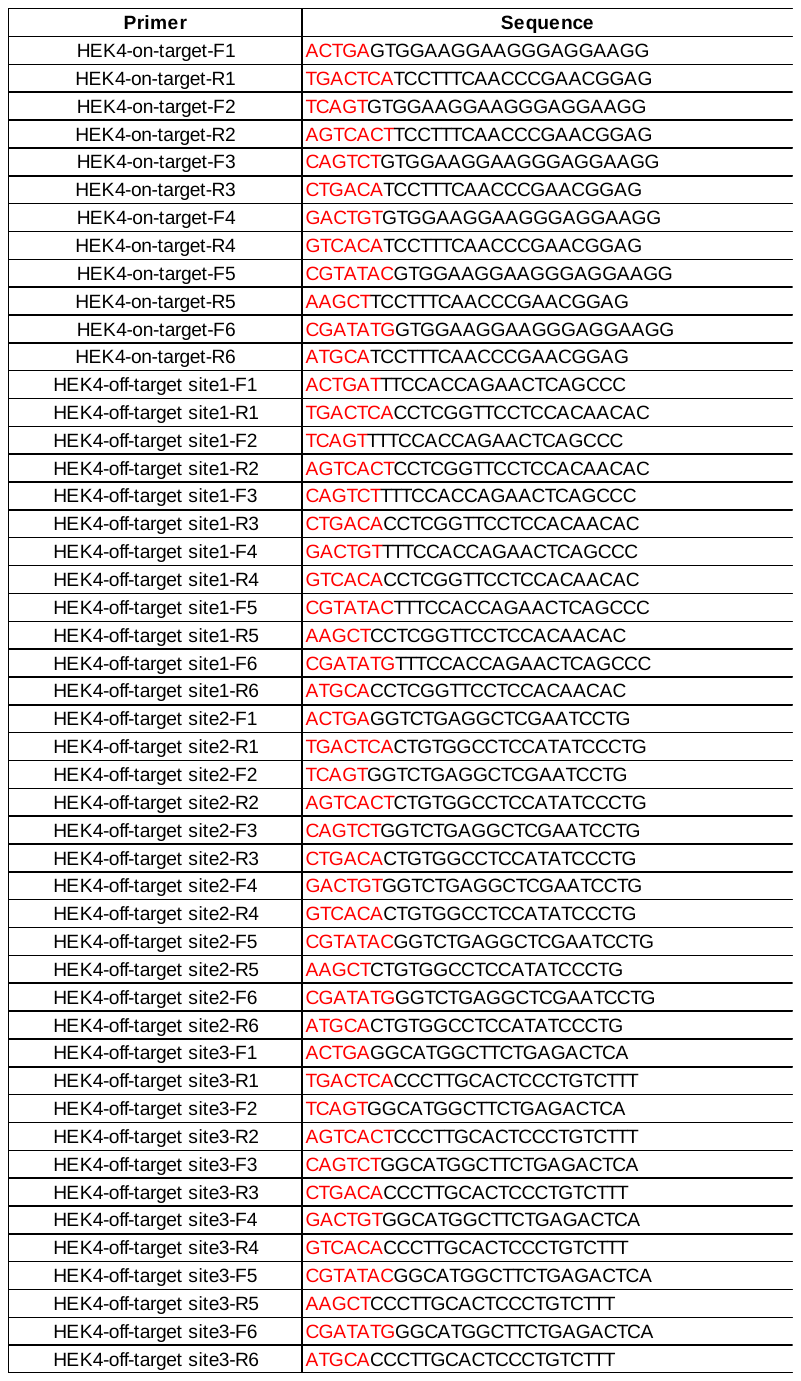

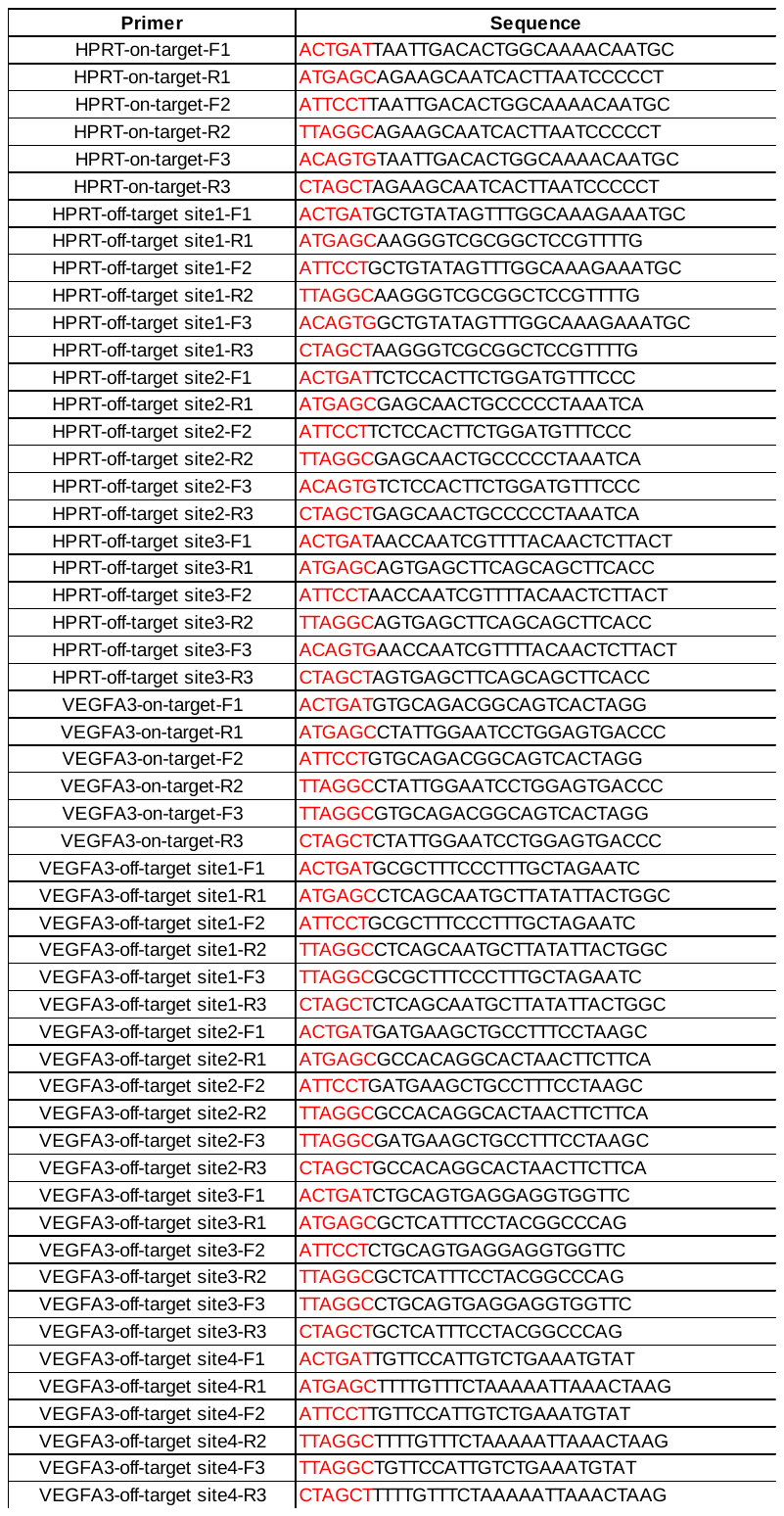
